# Tomtom-lite: Accelerating Tomtom enables large-scale and real-time motif similarity scoring

**DOI:** 10.1101/2025.05.27.656386

**Authors:** Jacob Schreiber

## Abstract

**Summary:** Pairwise sequence similarity is a core operation in genomic analysis, yet most attention has been given to sequences made up of discrete characters. With the growing prevalence of machine learning, calculating similarities for sequences of continuous representations, e.g. frequency-based position-weight matrices (PWMs), attribution-based contribution-weight matrices, and even learned embeddings, is taking on newfound importance. Tomtom has previously been proposed as an algorithm for identifying pairs of PWMs whose similarity is statistically significant, but the implementation remains inefficient for both real-time and large-scale analysis. Accordingly, we have re-implemented Tomtom as a numba-accelerated Python function that is natively multi-threaded, avoids cache misses, more efficiently caches intermediate values, and uses approximations at compute bottlenecks. Here, we provide a detailed description of the original Tomtom method (see Supplementary Note 1) and present results demonstrating that our re-implementation can achieve over a thousand-fold speedup compared with the original tool on reasonable tasks (see Supplementary Note 2).

**Availability and Implementation:** Our implementation of Tomtom is freely available as a Python package at https://github.com/jmschrei/memesuite-lite, which can be downloaded via pip install memelite.

## 1 Introduction

Calculating the similarity between a pair of sequences has been a central operation in genomics since its inception [1]. Initially, these algorithms assumed that both sequences would be discrete entities [2] (e.g., composed of nucleotides or amino acids), and were optimized given this constraint [3]. Yet, as the field expanded to the discovery of transcription factor (TF) binding sites and their regulatory effects, the motifs that were being discovered were not themselves discrete. Rather, because TFs do not need to bind to perfect matches, these motifs were encoded as position-weight matrices (PWMs) that were frequency-based, where each column encoded probabilities for each nucleotide appearing at each position [4]. New tools were necessary for learning PWMs from data [5, 6, 7], scanning these PWMs against discrete sequences [8, 9], and against each other. Tomtom emerged almost two decades ago as a popular algorithm for calculating similarities between a set of query PWMs and a set of target PWMs and subsequently converting these scores into p-values that account for the length and information content of the motif [10, 11].

The core conceptual innovation of Tomtom is the faithful calculation of null distributions. Briefly, the challenge is that one cannot assume the query and target PWMs are aligned or even the same length. Consequently, most PWM similarity calculations proceed by considering *all possible* ungapped alignments (and potentially reverse complements) and returning the *maximum* similarity score (Fig 1A). Because arriving at this score requires many operations, the null distributions for the statistical test must be calculated in a way that represents this entire series of operations. In their original work, Gupta *et al*. [10] propose Tomtom as an efficient algorithm for doing so, and a modification of Tomtom by Tanaka *et al*. [11] adjusts the similarity scores based on the number of unaligned positions. See Supplementary Note 1 for a complete description of the Tomtom algorithm, including implementation details.

**Figure 1.**
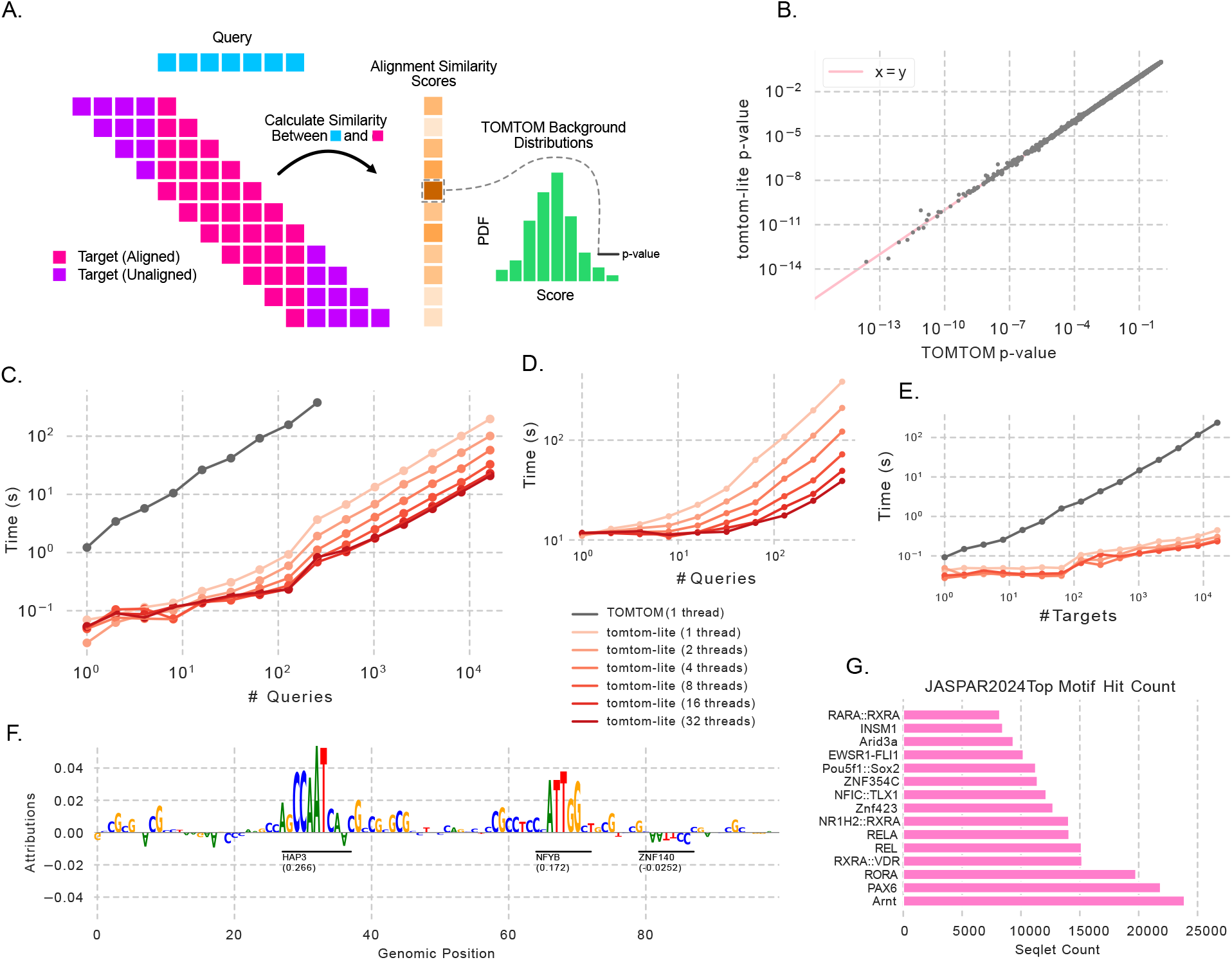
A schematic and results of our Tomtom implementation. (A) A schematic demonstrating that for each possible alignment (each row of purple/pink boxes) a score is calculated as the sum of all aligned columns between the target and the query (in pink), and the maximum score is kept and converted to a p-value. (B) P-values from Tomtom and tomtom-lite when comparing JASPAR2024 against itself. Only 1% of the *>*5M points are displayed. (C) Timings for Tomtom (in gray) and tomtom-lite with an increasing number of queries. (D) Timings when using a database of 1M random PWMs. (E) Timings when considering a target database of increasing size. (F) An example attribution track with seqlets called and annotated using Tomtom and JASPAR. (G) Count of the number of seqlets mapped to each motif in JASPAR according to tomtom-lite.

Being able to quickly calculate similarities between PWMs has recently taken on newfound importance as machine learning (ML) models have been integrated into genomics analyses. Two of the most prevalent forms of ML models have been supervised models that directly predict genomic activity [12, 13], and language models [14, 15] that capture the distribution of the genome [16, 17, 18, 19]. Although these models differ in significant ways, both have feature attribution methods that reveal which nucleotides drive model predictions, e.g. *in silico* saturation mutagenesis [20]. Spans of high-attribution nucleotides, called “seqlets”, can then be identified [21]. The computational challenge then becomes annotating these seqlets with the TF (or TF family) whose binding they most resemble. This can be done either by mapping the discrete sequence within seqlets to a motif database, or by mapping a PWM containing these attribution values (sometimes called a contribution-weight matrix or CWM [22]) to a database of attribution-based motifs. Tomtom is a natural solution to both problems.

Unfortunately, three practical challenges have limited Tomtom’s adoption. First, Tomtom does not scale well to large target databases despite being implemented in C. This is due, in large part, to the unnessesary re-calculation of null distributions for each query-target pair. Second, Tomtom is not natively multi-threaded and so requires additional command-line tools and expertise to utilize modern multicore computing systems. The third is that usage from an interactive environment or Jupyter notebook [23] requires writing the queries and targets out to disk, running the command-line tool, and then reading the results back from disk into memory. These steps can be burdensome and are error prone when trying to quickly do interactive analyses.

Our implementation of Tomtom (referred to hereafter as “tomtom-lite”) overcomes these challenges with a numba-accelerated Python function. Upon their first use, numba functions are compiled down to machine code that are similarly fast to their C counterparts [24]. These functions can use multi-threaded parallelism without significant code changes. Additionally, tomtom-lite caches null distributions across query-target pairs, organizes operations to avoid cache misses, uses an approximate median in a computational bottleneck, and allows for hashing of columns in the target database to reduce redundancy in the calculations. Together, these improvements make tomtom-lite up to three orders of magnitude faster than Tomtom in our evaluations.

As an initial check of our implementation, we compared the p-values produced by Tomtom and tomtom-lite when comparing the entire JASPAR2024 motif database [25] against itself. We observed a high correlation between their p-values (r=0.99981, Fig 1B), with tomtom-lite being ~417x faster than Tomtom (~2420s for Tomtom, ~5.8s for tomtom-lite). Importantly, the p-values have two regimes: *r* = 0.99981 when *p >* 1*e −* 5 containing 99.5% of comparisons, and *r* = 0.97 when *p ≤* 1*e −* 5, where the influence of the approximations is greater. These results suggest that the approximations have little effect when the goal is more discriminative in nature, e.g., identifying which targets might be good matches, but might influence results more when one cares about precisely ranking strong hits against each other.

We then evaluated how the two implementations scale with respect to various data properties. See Supplementary Note 2 for details on our exact evaluations. Both implementations scaled linearly with the number of queries but, due to differing scaling constants, tomtom-lite was 43x faster with 1 query and 265x faster with 128 queries (Fig 1C). Running tomtom-lite with 16,384 queries and 32 threads was faster than running Tomtom with 16 queries. We noticed that additional threads diminished in improvement past 8 and hypothesized that this was because the communication cost of managing that number of threads became higher than each task itself. Accordingly, we considered a more compute-intensive task where the target database was 1 million randomly generated PWMs that were longer, on average, than those in JASPAR. In this evaluation, we observed better scaling up to 32 threads (Figure 1D), suggesting that more threads will not always help when the target database is too small. Next, we found that both implementations scaled linearly with the number of targets but tomtom-lite scales significantly better because it caches null distributions. With only 1 target, tomtom-lite was only 2.1x faster with 1 thread and 2.89x faster with 8 threads, but with 16,384 targets tomtom-lite is 528x faster with 1 thread and 1,015x faster with 8 threads (Fig 1E).

We demonstrate these speed improvements in two practical settings: real-time interactive analysis, and large-scale motif similarity scoring. The first setting is invaluable for the exploration of data and model predictions, the generation of hypotheses, and the prototyping of new methods. The second setting is more representative of the computational burdens faced by analysis tools aiming to scale genome-wide or further. In both settings, we consider downstream applications of a ML model, ChromBPNet [12], that makes predictions for chromatin accessibility (specifically, ATAC-seq) in K562 from nucleotide sequence.

In our interactive example, we consider applying ChromBPNet to a specific locus of interest to identify motifs driving accessibility. Here, we follow the standard protocol for calculating DeepLIFT/SHAP attributions [26, 27] (see Supplementary Note 2 for details), identify seqlets, and then use tomtom-lite to annotate these seqlets using JASPAR2024. Our procedure identified 4 seqlets (Fig 1F, only 3 seqlets visualized), which tomtom-lite annotated in ~0.1 seconds and Tomtom annotated in ~2.5 seconds.

In our larger scale example, we consider the goal of counting the number of seqlets genome-wide that map to each motif in JASPAR2024. Specifically, we consider ChromBPNet attributions across all 203,804 ATAC-seq peaks in K562, which result in 634,776 seqlets. Annotating these seqlets using the JASPAR2024 database, which has 2,346 entries, takes only ~1,260 seconds (~20 minutes) using 8 threads. This speed makes it possible for one to perform genome-wide seqlet annotation on a commercially available laptop. When we consider the top motifs by count, we find that Arnt, PAX6, and RORA are in the top 15 motifs (Fig 1G).

Although our evaluations here focus on traditional PWMs where each column contains a value for each nucleotide or amino acid, Tomtom and tomtom-lite do not require this. Rather, they only requires that each column represent a position and that the features are semantically consistent across sequences. This means that one could natively apply Tomtom to the setting where PWMs contain embeddings of each nucleotide, such as those derived from a language model. Given the popularity of language models across domains and their demonstrated usefulness when applied to the genome, we anticipate that motif representations derived in this manner may augment frequency-or attribution-based PWMs.

Our results demonstrate that tomtom-lite is significantly faster than Tomtom and that these speed improvements have important practical benefits. These speed improvements are most beneficial at both ends of the compute spectrum: when the number of queries is small, potentially because one is doing interactive analyses, and when the number of queries and targets is very large. Given these benefits, we anticipate that tomtom-lite will be invaluable for tools involving genomic annotation and discovery.

## Acknowledgements

We thank Austin Wang, Justin Sanders, and Nikolaus Mandlburger for their feedback on the manuscript, and William Noble and Timothy Bailey for feedback and discussions on the original Tomtom implementation. Research at the Institute of Molecular Pathology (IMP) is supported by Boehringer Ingelheim GmbH and the Austrian Research Promotion Agency (FFG, FO999902549). For the purpose of Open Access, the authors have applied a CC BY public copyright license to any Author Accepted Manuscript (AAM) version arising from this submission. The computational results presented were obtained using the CLIP cluster (https://clip.science).

## Supplementary Note 1: Tomtom Algorithm

Tomtom is a statistical test that converts similarity scores between two motifs into p-values. A key challenge in calculating these p-values is that these motifs are not assumed to be aligned, and so the similarity is derived as the maximum similarity across all potential ungapped alignments (excluding insertions and deletions). Consequently, the primary contribution of Tomtom is the algorithm for faithfully calculating null distributions in a manner that accounts for this maximum operation across all possible alignments. Importantly, Tomtom is not itself a similarity score and so, conceptually, can be used with any similarity or distance score *S* between individual positions, though the original implementation only allows users to choose between a few hard-coded ones. In practice, additively decomposable score functions are significantly faster to compute and negative Euclidean distance works well. Here, we will provide a description for how to recreate the original Tomtom implementation, with notes on what tomtom-lite changed to improve speed.

Notationally, we will follow the original Tomtom work. 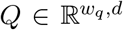 is a single “query” of length *w*_*q*_ and with an alphabet of size *d*. 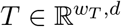 is the *concatenation* of all *n* sequences in a “target database” across the length dimension, with 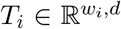 being the *i*-th entry in this database and 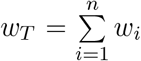. Here, the target database simply means a collection of sequences that are being scanned against the query, potentially a motif database like JASPAR2024. Finally, to be consistent with the original Tomtom work we will refer to each of the vectors of size *d* as “columns”, meaning that there are *w*_*q*_ columns in the query and *w*_*T*_ columns in the target database.

### 1.1 Integerized Similarity Matrix

The first step of Tomtom is to calculate a matrix of similarity scores between the columns in the query and those in the target database. We calculate 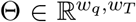 where each row represents one column in the query and each column represents one column in the target database and Θ_*i,j*_ = *S*(*Q*_*i*_, *T*^*j*^) where *T* ^*j*^ represents one column in the concatenation, as opposed to an entire target. As described by Tanaka *et al*., normalizing these scores accounts for differences in the number of unaligned columns when considering all potential alignments. This step is important to prevent single-column alignments on the edges that happen to match from scoring higher than an imperfect match between the core of the motifs. To do so, we calculate the median *m*_*i*_ for each of the *w*_*q*_ rows in Θ. In general, this normalization is robust to approximations of the median, and so tomtom-lite approximates this median for efficiency reasons (see Supplementary Note 2). Finally, these raw similarity scores are converted into integers ranging from 1 to *t* where *t* is the maximal similarity, which can be set by the user but defaults to 100. As a complete description, this procedure involves

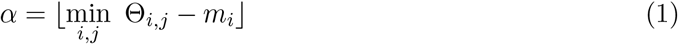

where *α* is the floor of the minimum median-normalized value in Θ,

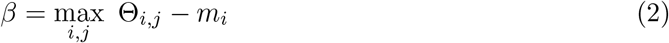

where *β* is the maximum median-normalized value (without taking the floor) in Θ, and

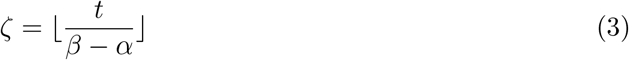

where *ζ* is the bin width. Given these values, we can directly convert the original (not median-normalized) similarity scores into integerized scores via

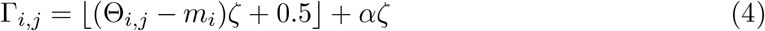

In the original Tomtom work by Gupta *et al*., Γ is used to represent both the initial matrix of continuous similarity scores and also the subsequent matrix of processed integer scores. For conceptual simplicity, we have separated them.

Admittedly, these steps are more complicated than traditional methods for converting a known range of number into integer bins, and it is unclear the value that they add, but they are necessary for perfectly reproducing the original Tomtom implementation.

#### 1.2 Null distributions

The next step is to calculate the null distributions of similarity scores to use in the statistical test. We use the plural because one distribution is needed for each target length from 1 to 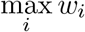, because differing target lengths means a different number of potential alignments we are taking the maximum operation over. We begin this step by calculating the marginal PDF of scores for each query column 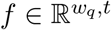. More formally,

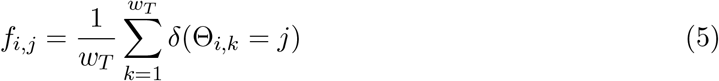

where *δ* is the Kronecker delta function which is 1 when the term is true and 0 otherwise. This corresponds to Eq. 2 in the original Tomtom paper and results in an *f* where the sum of each row is equal to 1 and the values are the fraction of entries in Γ_*i*_ that correspond to that integer value.

We then use these marginal PDFs to calculate PDFs for each *span* of query columns, with the crucial assumption necessary for calculating a null distribution that these marginal PDFs are independent from each other. These span score PDFs encode the probabilities of observing each integer score when the target is aligned to only this span of columns in the query, assuming independence between the query columns. They will be important for quickly calculating the null distributions because they will be used in the simulated alignment procedure. Because *T*_*i*_ can be shorter than *Q*, we must consider all internal spans in addition to those that originate or end at the edges of *Q*. Mathematically, each span score PDF is

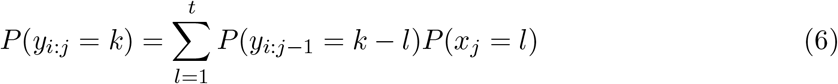

where *y*_*i*:*j*_ refers to the span PDF between columns *i* and *j* in the query and *x*_*j*_ refers to marginal distribution over the *j*-th column. Notably, *P* (*x*_*j*_) is already calculated in *f*_*j*_ and this equation can be solved recursively.

Accordingly, in our implementation we define 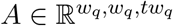 as the set of these span score PDFs. Here, the first dimension is the start of the span, the second dimension is the end of the span (inclusive), the third dimension is the size of the maximum integer score possible in any span, and the values are the probability of that score given that span. The lower triangle formed by the first two dimensions will be unfilled because spans cannot be negative. Because each *A*_*i,j*_ is the probability distribution *P* (*y*_*i*:*j*_), summing this tensor along the last dimension will produce a matrix where the upper triangle is entirely 1s.

Our first step in filling in *A* is to set the diagonals to be marginal distributions such that *A*_*i,i*_ = *f*_*i*_. Then, we proceed to fill out each row recursively as

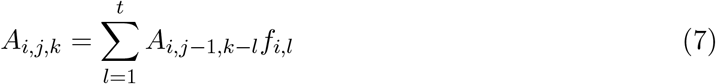

Essentially, the probability of observing the integer score *k* is the probability of observing *k − l* when the span were one column shorter multiplied by the probability of taking a step of size *l*, summed over all *l* between 1 and *t*. This is recursive because the distribution at index *j* can be directly calculated from the one at index *j −* 1.

Finally, we can use *A* to calculate our null distributions by simulating the alignment procedure. Because the returned score is the maximum of scores across all alignments, we need to calculate the distribution of the maximum integer that would be emitted by all relevant PDFs in *A*. Note that we are not simply taking the maximum probability at each position in the set of PDFs, but rather are calculating a new distribution describing the probability of the maximum sampled value across a set of PDFs being equal to a certain value. Because calculating this given a large set of distributions can be tricky to implement, we instead choose to iteratively calculate the maximum of a pair of distributions until the entire set has been exhausted. Specifically, we use

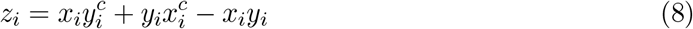

where *x*^*c*^ and *y*^*c*^ are the CDFs of *x* and *y* respectively and *z* is the PDF of the maximums.

Although applying Eq. 8 iteratively to a set of PDFs is straightforward, determining the set of PDFs to use requires some thought. Essentially, the number and composition of PDFs depends on the lengths of *Q* and *T*_*i*_. There are two situations. When *w*_*q*_ *≤ w*_*i*_, this set contains all PDFs in the first row of A except for the last one, 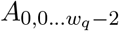, and all PDFs in the last column of A except the first one, 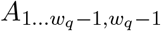 (the index 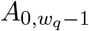 being excluded is the same in both cases, and is the top right index in the matrix). Then, this top-right index 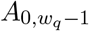 is included a total of *w*_*i*_ *− w*_*q*_ + 1 times; one time for each alignment of the *entire* query against a target that is longer than it. Because we are taking the max of a set of PDFs, including the same PDF multiple times does have an effect. When *w*_*q*_ *> w*_*i*_, we begin to use the internal spans. Specifically, if the canonical diagonal of a matrix is the 0-th diagonal, we use all PDFs in the *w*_*i*_ *−* 1st diagonal of *A*, all the PDFs in the first row leading up to the that diagonal, and all the PDFs in the final column leading to the bottom right index. For example, when *w*_*q*_ = 4 and *w*_*i*_ = 2, we would use *A*_0,0_, *A*_0,1_, *A*_1,2_, *A*_2,3_, *A*_3,3_.

The above process is repeated for each target length and results in a PDF. Because we want to quickly calculate p-values, we convert each PDF into 1 - CDF so that indexing into the distribution directly yields a p-value. Because our PDFs are over discrete integers, this can be achieved simply by taking the cumulative sum from the smallest to the largest integer in the distribution, and subtracting each value from 1.

A critical point is that these null distributions depend only on *w*_*i*_ and not on the composition of the target. In the original Tomtom implementation, this null distribution is recalculated for each pair of *Q* and *T*_*i*_, despite null distributions being identical for any target in a given target database of the same length. In tomtom-lite, we precalculate null distributions for all lengths from 1 to 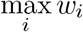 and re-use them across targets. Because calculating these distributions takes a significant amount of time, this change is the primary speed improvement in tomtom-lite.

### 1.3 Calculating p-values

At this point, all that is left is to calculate the similarity scores for each possible ungapped alignment between *Q* and *T*_*i*_, take the maximum, and convert this score into a p-value using the null distribution corresponding to the length of *T*_*i*_. Fortunately, this process is easy because we have already calculated the integerized distances between all columns in the query and all columns in the target database. Considering the slice of Γ corresponding only to the columns of *T*_*i*_, denoted *γ* (remember that Γ has *w*_*q*_ rows and *w*_*T*_ columns, spanning the entire target database, meaning *γ* has *w*_*q*_ rows and *w*_*i*_ columns), the similarity scores for the alignments can be easily calculated as the sum of the diagonals of *γ*. The maximal such diagonal-sum is then kept and converted to a p-value by indexing into the appropriate 1 - CDF. When considering reverse complements, the query and it’s reverse complement are both processed independently, the maximal score between the two is kept, and the p-value becomes 1 *−* (1 *− min*(*p*_1_, *p*_2_))^2^ where *p*_1_ and *p*_2_ are the relevant p-values.

### 1.4 Implementation Details for tomtom-lite

Despite tomtom-lite being a Python function, we have demonstrated that it is significantly faster than Tomtom, which is implemented in C. There are several factors that contribute to tomtom-lite’s speed, and they can be divided broadly into those that are beneficial without reducing precision, and those that are beneficial but do cause a loss of precision. In practice, we have found that the loss of precision for the default settings is minimal, but may become meaningful in non-standard settings or when changing the defaults.

#### 1.4.1 Speed Improvements Without Precision Loss

First, tomtom-lite is implemented using numba. Numba is a just-in-time (JIT) compiler that compiles functions into machine code the first time they are run. This offers C-like speed for numeric operations with Python-like syntax. Built-in caching functionality enables these compiled functions to be saved to disk, so the compilation cost only occurs the first time the function is run. Combined, this means that despite being available as a Python function, tomtom-lite would be as fast as a C implementation.

Second, numba enables multi-threaded parallelism without significant code changes and this parallelism is done across the processing of queries. Importantly, this is truly multi-threading with shared memory and not simply multi-processing, where multiple instances of Python are started and communication costs are high. This is possible because the compiled code is not bound by the global interpreter lock (GIL) that restrains Python code. Simply by using prange and using the `parallel=True` decorator appropriately, for loops can be converted into being parallel so long as the entire operation remains within numba (e.g., no calls to pure Python functions or globals) and calls within the loop do not depend on each other. This is satisfied in our implementation because each query can be processed independently. To avoid having each thread allocating small amounts of memory for each array needed for internal computation, tomtom-lite allocates a scratchpad before entering the loop whose first dimension is equal to the number of threads being used. Each thread uses the portion of the scratchpad allocated to its thread ID to avoid interfering with each other’s computations. The final output of each thread is written out to a pre-allocated results block.

Third, through careful implementation, tomtom-lite dramatically reduce the number of cache misses when calculating alignment scores. At a high level, when memory is accessed on modern systems, the elements surrounding a desired index are usually also loaded. Because the cost of starting an I/O operation is significantly higher than continuing it or performing compute on those elements, implementations that can make use of these adjacent elements can be several times faster than those that do not, even though, conceptually, the same set of operations are being performed. A “cache miss” is when elements are loaded because they are adjacent to earlier elements, then discarded, then directly accessed later on, and represent a wasted opportunity to eliminate an I/O call.

The potential for cache misses arises when summing the diagonals of *γ*. If we access an element and load the next elements in the row but then need to access the next element in the diagonal, we get an almost maximal number of cache misses because our operations are structured in a way to almost never make use of adjacent data. tomtom-lite eliminates these cache misses by, instead of proceeding along diagonals, proceeding along rows in the data and adding the element to an array of shape *w*_*q*_ +*w*_*i*_ *−*1. By ordering the operations in this manner, we observe a ~2-4x speed improvement in the step at the cost of having to keep the scores from all alignments in memory and taking the max at the end, instead of being able to keep a running maximum across diagonals.

#### 1.4.2 Speed Improvements with Precision Loss

In the process of turning Θ into Γ, the median of each row is subtracted from elements in that row. This has the effect of normalizing the score to account for the number of unaligned columns. However, calculating the exact median is time-consuming as doing so takes *O*(*w*_*T*_ *log w*_*T*_) time, which can grow to be quite large in some applications. Consequently, we replace this exact median with an approximate median that takes only *O*(*w*_*T*_ + *m*) time for a user-defined *m*, which is usually small with respect to *w*_*T*_. In this approximate formulation, the range between the minimum and maximum value (which is are calculated on-the-fly during the construction of Θ) is divided into *m* bins. After scanning through the *w*_*T*_ items in Θ_*i*_ to count the number of instances of each, we can scan through the *m* bins until we have encountered 50% of elements. For additional precision, we can store not only the count but the sum of elements within each bin and so, at the end, return the average value of elements within the bin. Although our empirical results have shown that Tomtom is robust to the use of an approximate median, the returned p-values may be different.

A final speed improvement involves reducing redundant calculations when calculating Θ. Essentially, many columns in *T* will be redundant due to the same character appearing in many motifs (e.g., a position in the PWM almost exclusively being an A, C, G, or T) and uninformative positions being fairly similar because they are almost the uniform distribution. Rather than re-calculating the similarity between the query column and these repeat columns in *T*, we can identify sets of similar columns in *T* as a preprocessing step and then calculate Θ over this set of non-redundant columns.

However, a key challenge is identifying these redundant columns in *T* quickly. The most conceptually straightforward way is to calculate the similarity between all columns in *T* and set a threshold for clustering, but this would likely take longer than the original task of calculating the similarity between *Q* and *T*. Accordingly, we use a hashing algorithm to identify similar columns. Briefly, for each of the *d* elements in *T*_*j*_, we divide the range between the minimum and maximum value into a user-defined number of bins *b*. Each column can then be assigned a hash index between 0 and *b*^*d*^ where elements with the same hash are similar, without needing to know anything more than the values within the column. Note that the number of hashes will likely far exceed the number of examples, but that we do not need to allocate memory for all *b*^*d*^ potential hashes.

When using hashed columns, minor algorithmic modifications need to be made to the other steps. When calculating the marginal distributions *f* we need to modify the Kronecker delta function to not return 1 when the statement is true, but rather return a count corresponding to the number of items in the hash index being considered. Once *f* is calculated, the null distributions can be calculated without modification.

A more complicated modification needs to be made when calculating the alignment scores while still minimizing cache misses. Specifically, instead of being able to extract *γ* directly as a series of contiguous columns, we have to look up the hash indexes for each column in *T*_*i*_ and iteratively load those columns. We proceed with a similar process to before, where the entirety of the column is used to update part of the vector of similarity scores. However, because we are operating on columns and the pre-loading mechanism works best on rows, we have to transpose the entire Γ matrix such that we are loading rows instead of columns. Because we are no longer loading a series of memory-adjacent rows (as there is no guarantee and it is actually quite unlikely that adjacent columns in *T*_*i*_ are adjacent in hash index) there is a small cost to doing it this way. However, despite slightly slowing down the calculation of similarity scores, this significantly speeds up the calculation of Γ to the point where it is no longer a compute bottleneck.

## Supplementary Note 2: Methods and Evaluations

In this work, we performed several experiments to demonstrate that tomtom-lite produces similar results to the original Tomtom implementation and to time the two implementations. The code and Jupyter notebooks for these comparisons can be found in the memesuite-lite GitHub repository: https://github.com/jmschrei/memesuite-lite. These comparisons were performed on a compute server with 76 Intel(R) Xeon(R) Gold 6138 CPU @ 2.00GHz cores. Many of these comparisons involved the JASPAR2024 motif database (called “JASPAR” for the rest of this note). We used the non-redundant core version, which can be be found at https://jaspar.elixir.no/download/data/2024/CORE/JASPAR2024_CORE_non-redundant_pfms_meme.txt

In these comparisons, each call to the Tomtom command-line tool was as follows:

`tomtom q.meme t.meme -motif-pseudo 0 -thresh 1 -text -verbosity 1 > o.tomtom`

where q.meme is a MEME-formatted file containing the queries, t.meme is a MEME-formatted file containing the targets, and o.tomtom is a TSV containing the outputs of Tomtom. Our timings for Tomtom include *only* running this command, and *do not* include the creation of q.meme and t.meme or subsequent reading of o.Tomtom into memory. Similarly, when running our Python function we only time the running of the function itself.

### 1.1.1 Comparing p-values between Tomtom and tomtom-lite

Our first evaluation involved comparing the p-values produced by Tomtom and by tomtom-lite to ensure that they were similar. This comparison was done by using JASPAR for both the queries and the targets. After running and timing the two commands, we constructed a p-value similarity matrix that was square because the number of queries and targets was the same. Because self-comparisons are definitionally very similar and because Tomtom uses 64-bit operations by default and tomtom-lite uses 32-bit operations by default, we set the diagonal of this matrix to 1 to reduce the influence of machine precision. We report the Pearson correlation

in two regimes: when the Tomtom p-values are above 1*e −* 5 and when they are below that.

### 1.1.2 Increasing Query Size

Our next evaluation was to consider timings when running an increasing number of queries against JASPAR. These queries are derived from the JASPAR itself to ensure a biologically plausible comparison. Since our comparisons ultimately end up considering many more queries than are in the JASPAR motif, we end up cycling through it multiple times. Each query is independent from the others, and so running multiple identical queries through a comparison does not influence our timing (or statistical) results.

Having shown that the benefits of multithreading diminish with increasing threads, we considered a second target database that is comprised entirely of random PWMs. These PWMs were generated using the following procedure: first, generate motifs of alphabet size 4 with width 15 by independently randomly drawing from *N* (0, 1), second, exponentiate these values to get strictly positive values, and third, divide each column by the sum of the columns to convert each value into something resembling a probability. A benefit of this exponentiation operation is that it results in columns that are strongly biased towards one character at each position, which is similar to real motifs. We then follow the same procedure for generating queries as when we used JASPAR, except without the need to cycle through the target database multiple times because it is so big.

### 1.1.3 Increase Target Size

We then turned to considering timings when the query size is held constant and the target size is fixed. Because multithreading was not shown to be particularly effecive when target databases were small, we considered only up to 8 threads and used only 20 queries. We constructed these target databases as the first *n* entries in JASPAR, looping back over the JASPAR once we have included all of the motifs, in a similar manner to how we constructed large query sets previously.

## 2 ChromBPNet and Attributions

Finally, we considered two evaluations involving feature attributions from a ChromBPNet model. We downloaded the ChromBPNet model for ATAC-seq signal in K562 from the ENCODE Project (https://www.encodeproject.org, accession ENCSR868FGK). This accession corresponds to an archive of five ChromBPNet models that have been trained and evaluated on different folds, and we use only the one that is trained on fold 0.. We use the bp-netlite repository (https://github.com/jmschrei/bpnet-lite) for loading this ChromBP-Net model into PyTorch and the tangermeme repository (https://github.com/jmschrei/tangermeme) for subsequent analyses, specifically, the implementation of DeepLIFT/SHAP, the recursive seqlet caller, the seqlet annotation function that uses tomtom-lite’s Tomtom implementation, and the attribution plotting utilities.

In the interactive example, we considered an enhancer of GATA2 whose coordinates are at chr3:128481848-128493691 on hg38. We extracted a 2,114 bp window centered at the middle of these coordinates and calculated feature attributions using DeepLIFT/SHAP with default settings (20 dinucleotide shuffles as the null distribution). To call seqlets, we used an unpublished recursive seqlet calling algorithm based on a statistical test and a recursive definition that each span within a seqlet must also be called as a seqlet. These seqlets were then annotated by taking the underlying discrete sequences and mapping them to JASPAR using tomtom-lite’s implementation of Tomtom.

In our larger-scale example, we repeated the above procedure across all ATAC-seq peaks in K562. We began by downloading the peak calls from ENCODE (accession ENCFF558BLC). We then calculated attributions for each peak and called seqlets using the exact same procedure as above except that we only used 5 dinucleotide shuffles for DeepLIFT/SHAP. This was for computational efficiency, as calculating attributions with even only 5 shuffles took several hours. Finally, we used tomtom-lite to map the identified seqlets to JASPAR using 8 threads.

Code reproducing these analyses can be found in the tutorials folder of the memesuite repository, in the notebook denoted as belonging to the application note.

## References

[1] TF Smith and MS Waterman. Identification of common molecular subsequences. J. Mol. Biol., 147(1):195–197, March 1981.

[2] Waqar Haque, Alex Aravind, and Bharath Reddy. Pairwise sequence alignment algorithms: a survey. In Proceedings of the 2009 conference on Information Science, Technology and Applications, New York, NY, USA, March 2009. ACM.

[3] F Chiaromonte, VB Yap, and W Miller. Scoring pairwise genomic sequence alignments. Pac. Symp. Biocomput., pages 115–126, 2002.

[4] Arttu Jolma, Jian Yan, Thomas Whitington, Jarkko Toivonen, Kazuhiro R Nitta, Pasi Rastas, Ekaterina Morgunova, Martin Enge, Mikko Taipale, Gonghong Wei, Kimmo Palin, Juan M Vaquerizas, Renaud Vincentelli, Nicholas M Luscombe, Timothy R Hughes, Patrick Lemaire, Esko Ukkonen, Teemu Kivioja, and Jussi Taipale. DNA-binding specificities of human transcription factors. Cell, 152(1-2):327–339, January 2013.

[5] TL Bailey and C Elkan. Fitting a mixture model by expectation maximization to discover motifs in biopolymers. Proc. Int. Conf. Intell. Syst. Mol. Biol., 2:28–36, 1994.

[6] Xiaotu Ma, Ashwinikumar Kulkarni, Zhihua Zhang, Zhenyu Xuan, Robert Serfling, and Michael Q Zhang. A highly efficient and effective motif discovery method for ChIPseq/ChIP-chip data using positional information. Nucleic Acids Res., 40(7):e50, April 2012.

[7] Timothy L Bailey. STREME: accurate and versatile sequence motif discovery. Bioinformatics, 37(18):2834–2840, September 2021.

[8] Janne Korhonen, Petri Martinmäki, Cinzia Pizzi, Pasi Rastas, and Esko Ukkonen. MOODS: fast search for position weight matrix matches in DNA sequences. Bioinformatics, 25(23):3181–3182, December 2009.

[9] Charles E Grant, Timothy L Bailey, and William Stafford Noble. FIMO: scanning for occurrences of a given motif. Bioinformatics, 27(7):1017–1018, April 2011.

[10] Shobhit Gupta, John A Stamatoyannopoulos, Timothy L Bailey, and William Stafford Noble. Quantifying similarity between motifs. Genome Biol., 8(2):R24, February 2007.

[11] Emi Tanaka, Timothy Bailey, Charles E Grant, William Stafford Noble, and Uri Keich. Improved similarity scores for comparing motifs. Bioinformatics, 27(12):1603–1609, June 2011.

[12] Anusri Pampari, Anna Shcherbina, Evgeny Z Kvon, Michael Kosicki, Surag Nair, Soumya Kundu, Arwa S Kathiria, Viviana I Risca, Kristiina Kuningas, Kaur Alasoo, William James Greenleaf, Len A Pennacchio, and Anshul Kundaje. ChromBPNet: bias factorized, base-resolution deep learning models of chromatin accessibility reveal cisregulatory sequence syntax, transcription factor footprints and regulatory variants. bioRxivorg, page 2024.12.25.630221, January 2025.

[13] Žiga Avsec, Vikram Agarwal, Daniel Visentin, Joseph R Ledsam, Agnieszka Grabska-Barwinska, Kyle R Taylor, Yannis Assael, John Jumper, Pushmeet Kohli, and David R Kelley. Effective gene expression prediction from sequence by integrating long-range interactions. Nat. Methods, 18(10):1196–1203, October 2021.

[14] Eric Nguyen, Michael Poli, Marjan Faizi, Armin Thomas, Callum Birch-Sykes, Michael Wornow, Aman Patel, Clayton Rabideau, Stefano Massaroli, Yoshua Bengio, Stefano Ermon, Stephen A Baccus, and Chris Ré. HyenaDNA: Long-range genomic sequence modeling at single nucleotide resolution. arXiv [cs.LG], June 2023.

[15] Hugo Dalla-Torre, Liam Gonzalez, Javier Mendoza-Revilla, Nicolas Lopez Carranza, Adam Henryk Grzywaczewski, Francesco Oteri, Christian Dallago, Evan Trop, Bernardo P de Almeida, Hassan Sirelkhatim, Guillaume Richard, Marcin Skwark, Karim Beguir, Marie Lopez, and Thomas Pierrot. Nucleotide transformer: building and evaluating robust foundation models for human genomics. Nat. Methods, pages 1–11, November 2024.

[16] Maxwell W Libbrecht and William Stafford Noble. Machine learning applications in genetics and genomics. Nat. Rev. Genet., 16(6):321–332, June 2015.

[17] Jacob Schreiber and Ritambhara Singh. Machine learning for profile prediction in genomics. Curr. Opin. Chem. Biol., 65:35–41, December 2021.

[18] Xuehai Hu, Alisdair R Fernie, and Jianbing Yan. Deep learning in regulatory genomics: from identification to design. Current Opinion in Biotechnology, 79:102887, February 2023.

[19] Gökcen Eraslan, Žiga Avsec, Julien Gagneur, and Fabian J Theis. Deep learning: new computational modelling techniques for genomics. Nat. Rev. Genet., 20(7):389–403, July 2019.

[20] Gherman Novakovsky, Nick Dexter, Maxwell W Libbrecht, Wyeth W Wasserman, and Sara Mostafavi. Obtaining genetics insights from deep learning via explainable artificial intelligence. Nat. Rev. Genet., 24(2):125–137, February 2023.

[21] Avanti Shrikumar, Katherine Tian, Žiga Avsec, Anna Shcherbina, Abhimanyu Banerjee, Mahfuza Sharmin, Surag Nair, and Anshul Kundaje. Technical note on transcription factor motif discovery from importance scores (TF-MoDISco) version 0.5.6.5. arXiv [cs.LG], October 2018.

[22] Žiga Avsec, Melanie Weilert, Avanti Shrikumar, Sabrina Krueger, Amr Alexandari, Khyati Dalal, Robin Fropf, Charles McAnany, Julien Gagneur, Anshul Kundaje, and Julia Zeitlinger. Base-resolution models of transcription-factor binding reveal soft motif syntax. Nat. Genet., 53(3):354–366, March 2021.

[23] Thomas Kluyver, Benjamin Ragan-Kelley, Fernando Pérez, Brian Granger, Matthias Bussonnier, Jonathan Frederic, Kyle Kelley, Jessica Hamrick, Jason Grout, Sylvain Corlay, Paul Ivanov, Damián Avila, Safia Abdalla, Carol Willing, and Jupyter development team. Jupyter notebooks - a publishing format for reproducible computational workflows. In Fernando Loizides and Birgit Scmidt, editors, Positioning and Power in Academic Publishing: Players, Agents and Agendas, pages 87–90, plNetherlands, 2016. IOS Press.

[24] Siu Kwan Lam, Antoine Pitrou, and Stanley Seibert. Numba: a LLVM-based python JIT compiler. In Proceedings of the Second Workshop on the LLVM Compiler Infrastructure in HPC, New York, NY, USA, November 2015. ACM.

[25] Ieva Rauluseviciute, Rafael Riudavets-Puig, Romain Blanc-Mathieu, Jaime A Castro-Mondragon, Katalin Ferenc, Vipin Kumar, Roza Berhanu Lemma, Jérémy Lucas, Jeanne Chéneby, Damir Baranasic, Aziz Khan, Oriol Fornes, Sveinung Gundersen, Morten Johansen, Eivind Hovig, Boris Lenhard, Albin Sandelin, Wyeth W Wasserman, François Parcy, and Anthony Mathelier. JASPAR 2024: 20th anniversary of the open-access database of transcription factor binding profiles. Nucleic Acids Res., 52(D1):D174–D182, January 2024.

[26] Avanti Shrikumar, Peyton Greenside, and Anshul Kundaje. Learning important features through propagating activation differences. arXiv [cs.CV], April 2017.

[27] Scott Lundberg and Su-In Lee. A unified approach to interpreting model predictions. arXiv [cs.AI], May 2017.

